# Homologous Recombination as an Evolutionary Force in African Swine Fever Viruses

**DOI:** 10.1101/460832

**Authors:** Zhaozhong Zhu, Chao-Ting Xiao, Yunshi Fan, Zena Cai, Congyu Lu, Gaihua Zhang, Taijiao Jiang, Yongjun Tan, Yousong Peng

## Abstract

Recent outbreaks of African swine fever virus (ASFV) in China severely influenced the swine industry of the country. Currently, there is no effective vaccine or drugs against ASFVs. How to effectively control the virus is challenging. In this study, we have analyzed all the publicly available ASFV genomes and demonstrated that there was a large genetic diversity of ASFV genomes. Interestingly, the genetic diversity was mainly caused by extensive genomic insertions and/or deletions (indels) instead of the point mutations. The genomic diversity of the virus resulted in proteome diversity. Over 250 types of proteins were inferred from the ASFV genomes, among which only 144 were observed in all analyzed viruses. Further analyses showed that the homologous recombination may contribute much to the indels, as supported by significant associations between the occurrence of extensive recombination events and the indels in the ASFV genomes. Repeated elements of dozens of nucleotides in length were observed to widely distribute and cluster in the adjacent positions of ASFV genomes, which may facilitate the occurrence of homologous recombination. Moreover, two enzymes, which were possibly related to the homologous recombination, i.e., a Lambda-like exonuclease with a YqaJ-like viral recombinase domain, and a DNA topoisomerase II, were found to be conservative in all the analyzed ASFVs. This work highlighted the importance of the homologous recombination in the evolution of the ASFVs, and helped with the strategy development of the prevention and control of the virus.

## Introduction

African swine fever virus (ASFV), the causative agent of African swine fever (ASF), is a complex, large, icosahedral multi-enveloped DNA virus. It is classified as the only member in the family *Asfarviridae* ^1,2^. The genome of the virus belongs to double-stranded DNA, with the size ranging from 170 kb to 190 kb ^3^. ASFV mainly infect suids and soft ticks. The suids include domestic pigs and wild boars, and were reported as the natural hosts of the virus ^4,5^. ASFV was firstly discovered in Kenya in 1921 ^6^. It remained restricted in Africa till 1957, when it was reported in Spain and Portugal. Up to now, the virus has caused ASF outbreaks in more than fifty countries in Africa, Europe, Asia, and South America ^4^. The latest reports showed that the virus has caused outbreaks in more than fifteen provinces in China ^7,8^. Because of the high lethality of ASFV in domestic pigs, the most commonly used strategies to control the virus were the massive culling campaigns and the restriction of pig movement ^5^. Both strategies have resulted in a huge economic loss for pig industry and affected people’s livelihoods. Unfortunately, currently there is no available effective vaccine against ASFVs.

Many efforts have been devoted to developing the vaccine for the ASFV ^1,5,9-11^, however, most of these attempts failed. One of the most important reasons was the complex composition of the antigenic proteins ^5,12^. Previous reports showed that p72, p30, and p54 were the three important antigenic proteins during the infection of ASFVs, but the immunity against them could only provide a partial protection ^12,13^. Many other proteins or other factors such as phospholipid composition may also influence the antigen of the virus ^12^. Therefore, it is necessary to understand the mechanisms of the antigen diversity of the ASFV virus ^1^.

The genetic diversity of ASFVs has been investigated in many studies. The ASFV genome encodes over 150 proteins, including viral enzymes, viral transcription and replication-related proteins, structural proteins, other proteins involved in the virus assembly, the evading of host defense systems and the modulation of host cell function, etc ^3,14,15^. For example, the transcription of the virus is independent on the host RNA polymerase because the virus contains relevant enzymes and factors ^3^. The viral genome contains a conservative central region of about 125 kb and two variable ends, which results in the variable size of the genome ^3,16,17^. There are significant variations among the ASFV genomes due to the genomic insertion or deletion, such as the deletion of the multigene family (MGF) members ^3^. Although much progress have been made on genetic diversity of the virus, the extent and mechanisms are still not clear. Besides, most of these studies either only investigated the genetic diversity of some common genes, such as p72 and p54 ^18,19^, or only used one or several isolate genomes ^3,16,17^. The number of discovered viral genomes has increased rapidly as the development of DNA sequencing technology. Therefore, a comprehensive study on the genetic diversity of ASFVs is necessary.

Homologous recombination, which has been reported to occur in several groups of viruses ^20-23^, such as herpesvirus, retroviruses, and coronaviruses, has played an important role in viral evolution ^21^. A few studies on several ASFV genes have suggested the occurrence of homologous recombination in the evolution of ASFVs ^3,18^. However, a comprehensive study on the homologous recombination in ASFV at the genomic scale is lacking, and the role of the recombination on the genetic diversity and the evolution of the virus is still unknown. In this study, we have systematically investigated the genomic diversity and the homologous recombination of ASFVs based on the analysis on all the publicly available ASFV genomes. The results demonstrated that the homologous recombination contributed much to the genetic diversity of ASFVs. This work would help to understand the evolution of the ASFV and thus facilitate the prevention and control of the virus.

## Results

### 1 ASFV genomes

A total of 36 genome sequences of ASFVs were obtained from the NCBI GenBank database, which were listed in Table S1. They were mainly isolated from Africa and Europe during the years from 1950 to 2017. The size of the ASFV genomes ranged from 170,101 bp to 193,886 bp, averaged at 185,800 bp. The viral isolate Kenya50 had the largest size, while the isolate BA71V had the smallest size. No increasing or decreasing trend in the genome size was observed from 1950 to 2017 (Figure 1A), suggesting the dynamic changes of the viral genomes.

**Figure 1.**
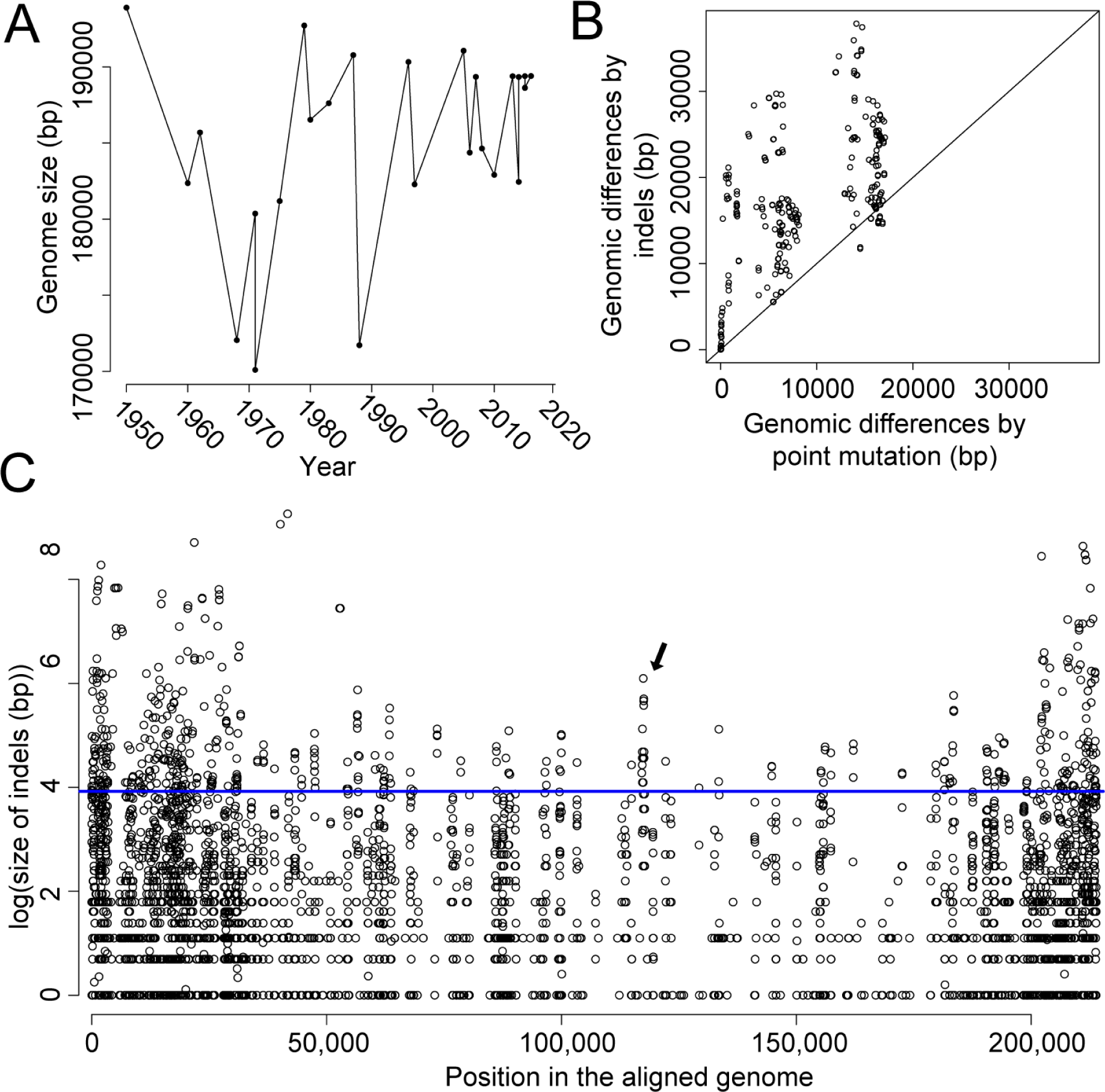
Genomic differences between ASFV genomes. (A) The variation of genome size along the isolation time of the virus. (B) Genomic differences caused by point mutations and indels. (C) The size and location of indels along the aligned genomes. For clarity, the natural logarithm of the indel size was used. Position for an indel is defined as the middle position of the indel. The average size was used if more than one indel was found in the position. The blue line refers to indel size of 50 bp.

### 2 Genomic diversity of ASFVs

Pairwise comparisons between ASFV genomes were conducted after the genome alignment. The average genomic difference between viruses was 24,570 bp, which accounts for more than 10% of the genome. Interestingly, the genomic differences caused by the insertions and deletions (indels) were much more significant than those caused by the point mutations (Figure 1B & Figure S1) in most cases. For example, there were 31,833 bp differences between virus Mkuzi79 and BA71V, 78% of which were caused by indels.

The size and position of indels in ASFV genomes were also analyzed. 70% of indels were no longer than 10 bp, and about 10% of indels were 50 bp or longer (Figure S2). The occurrence of indels was much more frequent in both ends of the genome, especially in the 5’ end (Figure 1C). Besides, the size of indels in both ends was also much larger than that in the middle region. Large indels with over 50 bp (above the blue line in Figure 1C) were mostly observed in both ends. It should be noted that the variation to some extent was observed in the middle region (marked in black arrow), which were considered to be conservative in previous reports.

### 3 Proteome diversity of ASFV

Genomic diversity could lead to proteome diversity. Therefore, the proteome diversity of ASFVs was further analyzed. Firstly, the candidate proteins encoded by ASFV genomes were inferred (Materials and Methods) (Table S2). The plus strand encoded 95-126 proteins, with an average of 109 proteins; the minus strand encoded 106-128 proteins, with an average of 118 proteins. Considering both the plus and minus strands, the ASFV genome encoded 205-254 proteins, with an average of 227 proteins. The viral isolate Russia14 encoded most proteins, although the size of this genome was not the largest. The ratio of coding region in each genome ranged from 88% to 91%.

Furthermore, the ortholog or paralog groups based on sequence homology were identified. A total of 252 protein groups plus 28 singletons were obtained (Table S3), each of which stood for one type of protein encoded by ASFV genomes. The obtained proteins contained almost all the proteins identified in previous experiments (Table S3). Each protein group included 2-99 proteins. Only 144 protein groups were observed in all 36 ASFV viruses, which could be considered as core protein sets of the virus, and were mainly encoded by both plus and minus strands in the middle regions (Figure 2). The protein groups could be further separated into seven classes by function based on previous studies (Figures 2 & S3). Only about 30% of protein groups were observed to have the known functions, including replication and transcription (in red), host cell interactions (in magenta), structure and morphogenesis (in blue), and enzymes (in yellow). Most of the above-described protein groups with the known functions belonged to the core proteins of the virus. In addition, forty protein groups belonged to “Multigene Families (MGF)” (in cyan), most of which had unknown functions. The MGFs were encoded by both ends of the genome. Besides, the remaining 146 protein groups belonged to either the class of “Proteins with unknown function” (in gray) or “Hypothetical proteins” (in black).

**Figure 2.**
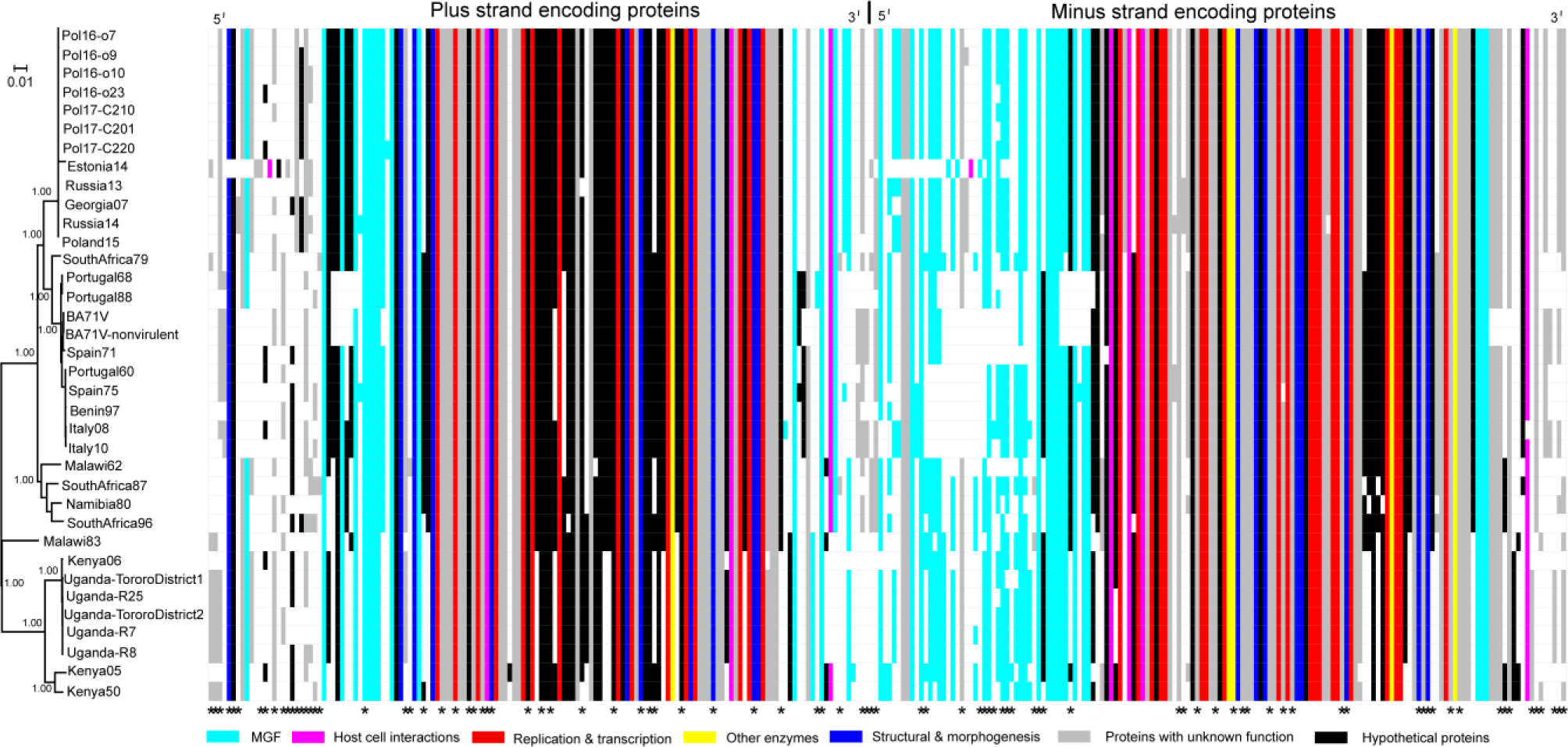
The phylogenetic tree of ASFVs and the alignment of their proteomes in plus (left side) and minus strand (right side). Each row refers to the proteome of the virus in the phylogenetic tree; each column refers to one protein group. Protein groups were colored according to their functions. “White” refers to no protein group in the virus. Asterisks in the bottom refer to membrane proteins. For clarity, the singletons were ignored in the alignment.

Analysis of the protein conservation showed that except the functional class of MGFs, the proteins in other functional classes had an average of pairwise sequence identities greater than 90% (Figure S4). The proteins in functional classes of “Other enzymes” and “Replication & transcription” were most conservative, with average pairwise sequence identities larger than 95%. Proteins of these two functional classes also had the smallest ratios of dN/dS (Figure S5), suggesting strong negative selection on them. While the proteins in the functional class of MGF and “Hypothetical proteins” had the largest ratio of dN/dS. The hypothetical proteins had a median dN/dS ratio of 0.82, suggesting strong positive selection on these proteins.

Membrane proteins, which may be located in the inner or outer envelope, were observed to be distributed widely in the proteome of ASFVs (marked with asterisks in Figure 2). A total of 67 protein groups belonged to membrane proteins, including 35 in the core protein groups, such as p54 and EP402R. Among the membrane proteins, only 11 protein groups had the known functions, including 8 protein groups in the functional class of “Structural and morphogenesis”, and 1 protein group in each functional class of “Host cell interactions”, “Replication & transcription” and “Other enzymes”.

Thirty-one protein groups were observed to have paralogs (duplicated proteins) in at least one virus (colored in Figure S6). They were mostly located in both ends of the genome. Thirteen of them belonged to MGFs. In addition, two protein groups, “DP71L” and “DP96R”, belonged to the class of “Host cell interactions”. The rest protein groups belonged to either the class of “Proteins with unknown function” or “Hypothetical proteins”. Most of the paralogs were clustered in adjacent positions. Exceptions were observed for some protein groups which were encoded by the first one to three thousands nucleotides in the plus and minus strands, such as the protein group “p01990-3L” (marked with black arrows in Figure S6). Further analysis showed that a segment of 200-3000 bp was exactly the same in the beginning of the plus and minus strands in most viral genomes (Table S4).

Extensive insertion and deletions of proteins were observed in the proteome of ASFVs after alignment. Viruses in the adjacent positions in the phylogenetic tree tended to have similar proteomes. The number of different proteins between different viruses ranged from 1 to 84, with an average of 43, which was about one-fifth of the viral proteome. The differences of the proteome among the viruses were mainly caused by proteins of the class of “Hypothetical proteins”, “Proteins with unknown function” and “MGF” (Figure 2).

### 4 Extensive homologous recombination in ASFV genomes

As numerous indels have been revealed in the ASFV genomes, then, we investigated the mechanism of generating indels. According to the results in previous studies, three factors may contribute to the extensive indels in ASFVs: replication slippage, retrotransposition and recombination ^23^. Replication slippage mainly produced duplications of short genetic sequences and may cause short indels, but it is unlikely to generate large indels observed in ASFVs. Retrotransposition can result in duplication of large genetic sequences or genes, but the location of duplicates would be randomly distributed in the genome. However, most paralogs shown in Figure S6 were clustered in adjacent positions, thus these paralogs may be not caused by retrotransposition. Besides, no retrotransposons were observed in the analyzed ASFV genomes (as described in Materials and Methods).

Finally, we investigated the role of recombination in the generation of indels in the ASFV genomes. The analyses on the recombination showed that there were a total of 103 recombination events, and each ASFV genome had 3-22 recombination events (Figure 3 & Table S5). The virus isolate SouthAfrica79 experienced the largest number of recombination events. On average, each virus experienced 11 recombination events. The sizes of recombination region ranged from 174 to 22,628 bp. The ratio of recombination region in each genome ranged from 2% to 27%. In total, the regions in the ASFV genomes involved in all recombination events covered a total of 101,569 nucleotide positions, accounting for 47% of the aligned genome. Most recombination events happened at both ends, especially at the 5’ end. Interestingly, the recombination event in the aligned genomes was observed to be consistent with the ratio of the gap in the genome (the bottom of Figure 3). Almost all the recombination events happened in or close to the gap-rich regions where the indels were observed. The ratios of the gaps in the recombination regions were found to be much larger than those in other regions (Figure S7). Further comparison of the number of indels in the recombination regions and other regions showed that for indels of varing length, such as those greater than 5, 10, or 50 bp, the number of indels in the recombination regions was much larger than those in other regions (Figure S8).

**Figure 3.**
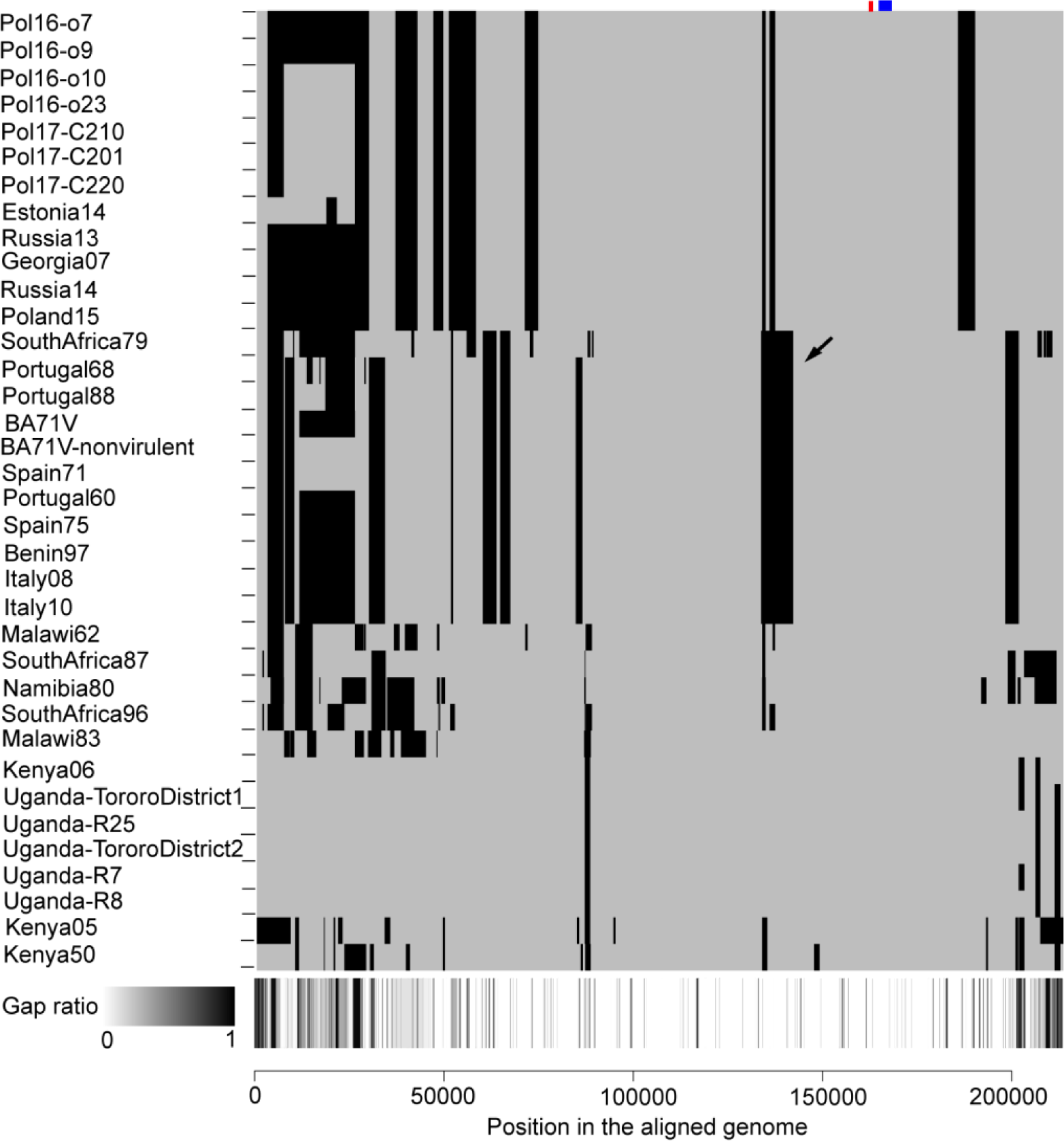
Recombination of ASFV genomes. The black areas indicate the recombination region for each genome. The bottom panel shows the ratio of gap in each position of the aligned genome. The panel uses the grayscale color bar at the bottom-left. The red and blue rectangles in the top-right indicate the coding region of pD345L (Lambda-like exonuclease) and P1192R (DNA topoisomerase II), respectively. The black arrow refers to the recombination event displayed in Figure 4.

Figure 4 illustrates the recombination event in 11 viral isolates (colored in red), including two viral isolates from Africa (SouthAfrica79 and Benin97) and nine viral isolates from Europe. These 11 viral isolates formed a separate lineage in the phylogenetic tree. The recombination region ranged from 133,683 to 142,222 bp, located in the central conservative region of the genome (shown by the black arrow in Figure 3). In the phylogenetic tree built with genomic sequences without the recombination regions, the recombinants are the neighbors of a clade containing viruses from Eastern Europe countries (Figure 4A); while in the tree built with genomic sequences of the recombination regions, the recombinants are the descendants of Malawi62 from Africa (Figure 4B).

**Figure 4.**
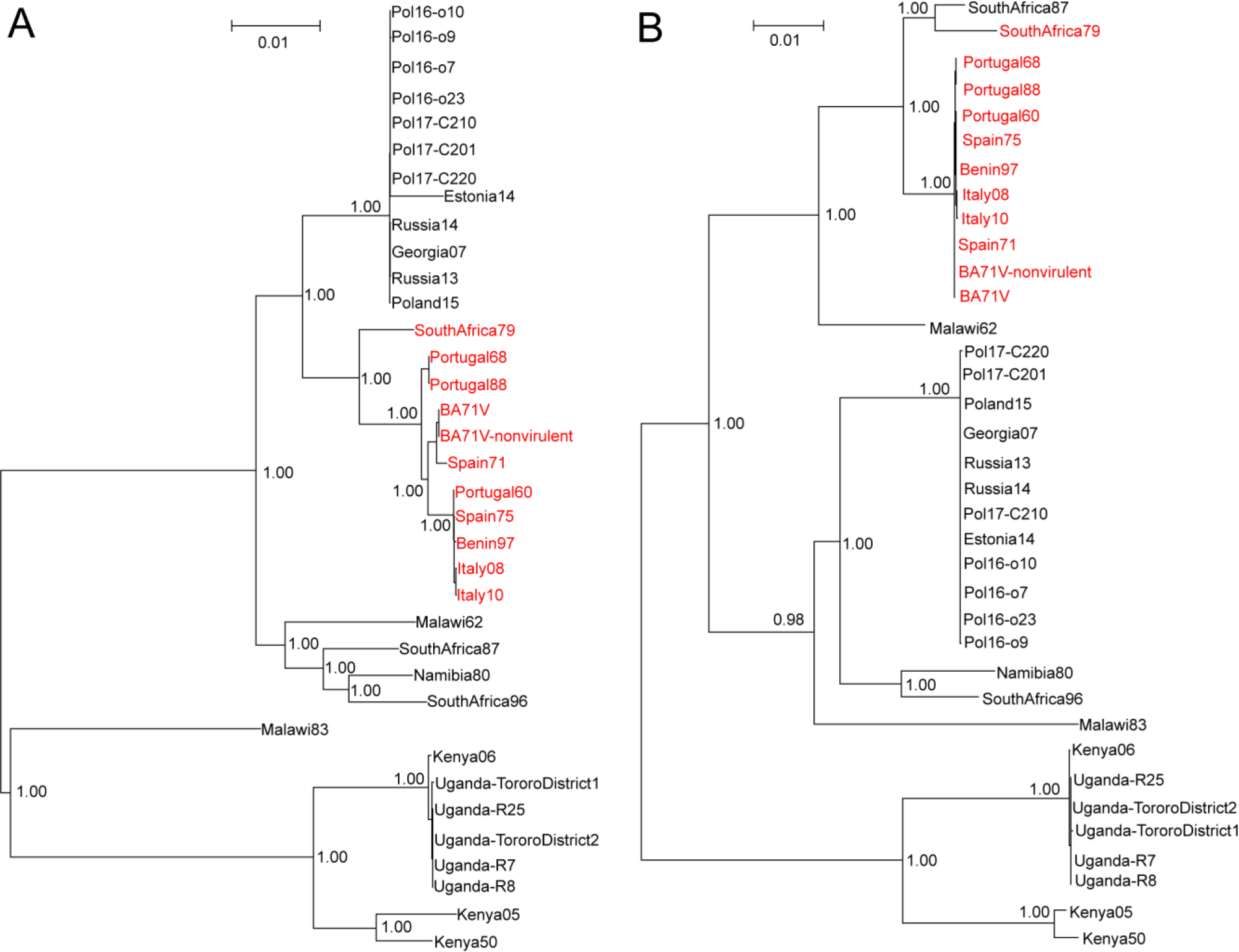
An example of recombination events in 11 ASFVs (colored in red). Figure (A) refers to the maximum-likelihood phylogenetic tree built with genome sequences without the recombination region (133,683-142,222 in the aligned genome). Figure (B) refers to the phylogenetic tree built with genome sequences of the recombination region. The numbers refer to the bootstrap values of nodes in the bootstrapping test with 100 replicates.

In addition, the indels introduced directly by the recombination were investigated. The results of comparing the sequence in the recombination regions of recombinants to that in the major parent viruses showed that an average of 37% of differences were caused by the indels in all the recombination events. Further comparison on the proteins showed that in 58 of 103 recombination events, there was at least one different protein encoded by the recombination region of the recombinants from that encoded by the major parent viruses.

Furthermore, the proteins involved in the recombination events were further analyzed. A total of 110 protein groups were involved in the recombination events, including 47 core protein groups and 63 variable protein groups (Table S3). 34 of 110 protein groups belonged to membrane proteins. Four protein groups of “Host cell interactions” (EP153R, A238L, DP96R and DP71L), eight protein groups of “Structure & morphogenesis” and nine protein groups of “Replication & transcription” were involved in the recombination events.

### 5 Identification of possible recombinase and DNA topoisomerase in ASFVs

Interestingly, we found a protein, named pD345L, denoted as the Lambda-like exonuclease, was possibly involved in the recombination because it contained the YqaJ-like viral recombinase domain. The protein pD345L has 345 amino acids and is encoded by the minus strand. It was highly conservative in ASFVs, with an average sequence identity of 96.7% between ASFVs. Even higher level of conservation was observed in the recombinase domain of pD345L, with an average sequence identify of 97.2%. Although the YqaJ-like recombinase domain was extensively distributed in Bacteria, Virus and Eukaryota, it was considered as the viral origin ^24^. The recombinase domain in ASFV was most similar to that in two giant viruses, Pacmanvirus and Kaumoebavirus (see Materials and Methods) which were possibly distant relatives of ASFVs ^25,26^.

Besides, topoisomerase was also reported to be related to homologous recombination. We found that a type II DNA topoisomerase, i.e., P1192R, exists in all analyzed ASFV isolates. P1192R was a protein including 1192 amino acids, and was encoded by the plus strand. P1192R is also conservative in these viruses, with an average sequence identity of 98.3% between ASFVs. Although pD345L and P1192R were encoded by different strands, they were encoded by the genomic sequences in adjacent regions: the former was encoded by the sequences in the positions of 162,236-163,273 (colored in red in Figure 3), while the latter was encoded by the sequences in the positions of 164,743-168,319 (colored in blue in Figure 3). Both enzymes were not involved in any recombination events. The phylogenetic trees for proteins pD345L and P1192R were similar to the tree built with the whole genome (Figure S9).

### 6 An abundance of repeated elements in ASFV genomes

Repeated elements could facilitate the homologous recombination. In this study, lots of repeated elements ranging from 5-100 bp were identified, and then the distribution of the repeated elements in the ASFV genomes was analyzed. As shown in Figure S10, the number of repeated elements in ASFV genomes decreased monotonously as the size of elements increased. Then, the distances between adjacent elements for a given repeated element was investigated (Figure 5A). As the size of the elements increased from 5 to 10, the average distance between the adjacent elements also increased because the number of repeated elements in the genome decreased. Interestingly, the average distance decreased as the size of the elements increased from 11 to 23; it reached to the minimum (136 bp) when the size was 23; then the distance kept unchanged as the size increased from 23 to 46; finally, it increased as the size of repeated element increased from 47 to 100. It should be noted that the average distance was still less than 400 bp even for the repeated elements of 100 bp. These phenomena suggested that the repeated elements of 11 bp or larger tended to cluster in the genome, especially for those of 23-46 bp.

**Figure 5.**
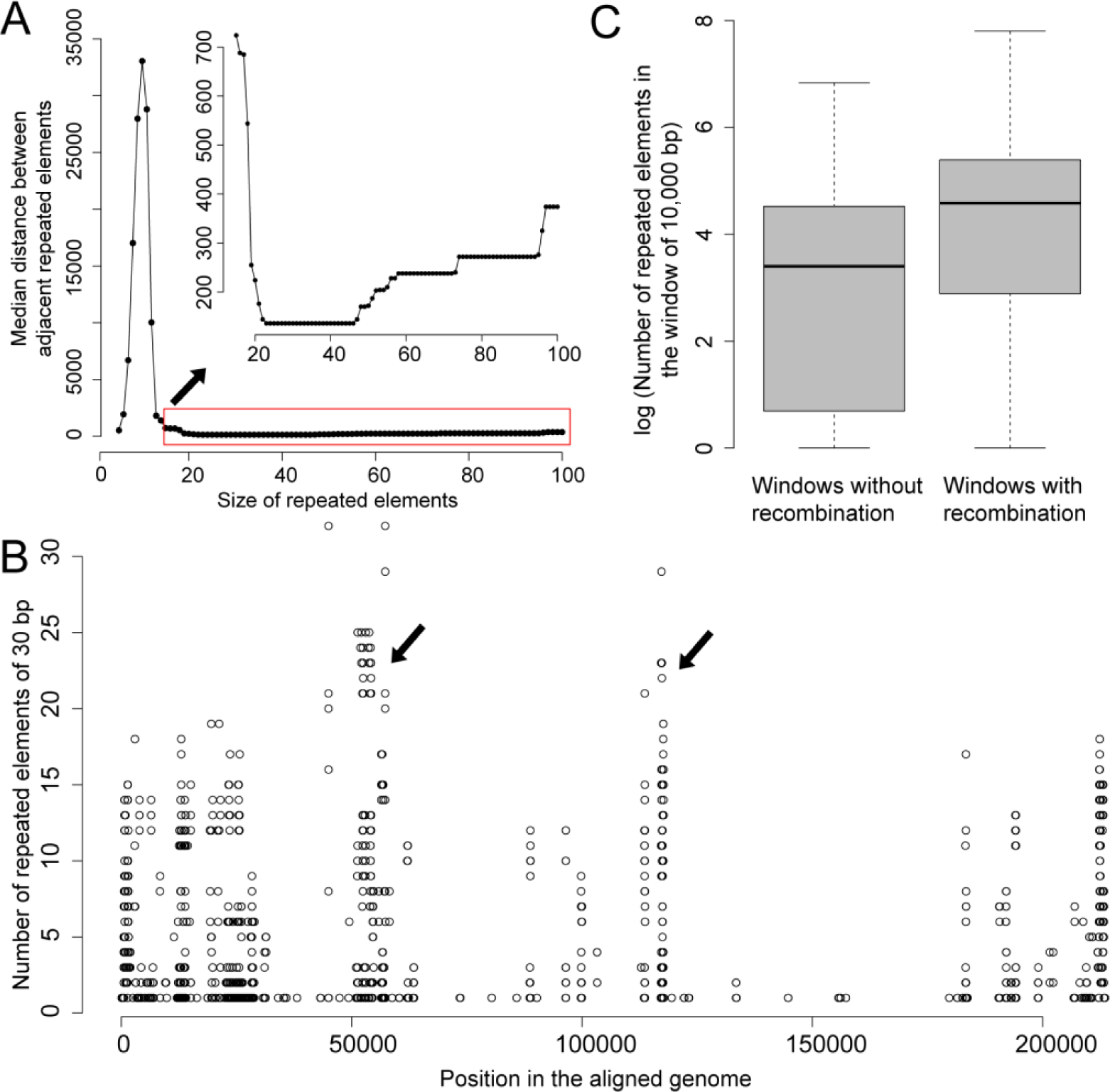
Distribution of the repeated elements. (A)The median distance between adjacent repeated elements versus the size of repeated elements. (B) Number of the repeated elements with the size of 30 bp observed in each genomic position. (C) Comparison of the number of repeated elements (15 bp in length) in the window of 10,000 bp with and without the recombination in viral genomes. For clarity, the natural logarithm of the number of repeated elements was used.

For example, when the size of elements was 30 bp, each genome had a median of 427 types of elements which repeated at least two times in the genome. Some elements appeared for over ten times in the genome, such as the element “AGGCGTTAAACATTAAAATTATTACTACTG” in the viral strain BA71V. The region covered by repeated elements accounted for 1%-3% of the genome in ASFVs. The distance between repeated elements was analyzed and demonstrated to have a median distance of 136 bp, suggesting they tend to cluster in adjacent regions. Figure 5B shows the distribution of repeated elements in the aligned genome. Most repeated elements were located at both ends of the genome. Besides, there were two clusters of repeated elements in the positions of around 55,000 bp and 120,000 bp (marked by black arrows), respectively.

Finally, the contribution of repeated elements to the recombination was investigated. For elements of 10 or more nucleotides, the number of repeated elements in the windows (2000-10,000bp in length) including the recombination was significantly larger than those without the recombination (Table S6). Figure 5C shows the comparison of the number of repeated elements (15 bp in length) in the windows of 10,000 bp with and without the recombination in viral genomes. The windows including the recombination had a mean of 194 repeated elements, which was twice of that in the windows without the recombination.

## Discussion

This work systematically analyzed the genetic diversity of ASFVs. The large genome of the virus enabled the encoding of an abundance of proteins. Each of the functions of the virus in its life circle could be accomplished by multiple proteins. For example, 35, 19, and 7 protein groups were involved in DNA replication and transcription, structure and morphogenesis, and host cell interactions, respectively. On one hand, this multiple protein mechanism could facilitate the efficient control of the host cell by protein-protein interactions, such as inhibiting the transcriptional activation of host immunomodulatory by A238L ^3^ and inhibiting Toll-like receptor 3 signaling pathways by I329L ^27^; on the other hand, this mechanism could facilitate the precise regulation of the viral activities. For example, the ASFV virus was considered to contain all the enzymes and factors which were required for the transcription and post-treatment of mRNAs ^3^.

Significant differences were observed among the different proteomes of ASFVs, which may be caused by the following two reasons: i) over 40% of the proteins were non-essential among ASFVs, and ASFVs may have variable number of these proteins; ii) there were 31 genes with replications in ASFV genomes. Diverse proteome among ASFVs may lead to diverse phenotype, such as the diversity in antigen and virulence. The diversity may result in a great challenge for the prevention and control of the virus. For example, the viruses with diverse antigens may need multiple types of vaccines because the effectivity of the cross-protection on viruses may be limited.

Although lots of efforts have been devoted to developing vaccines against ASFVs ^1,10-12^, unfortunately, most of the attempts have been unsuccessful. The failure could be caused by many factors ^12^, including the absence of neutralizing antibodies, the diverse antigen-related proteins, the complexity of neutralization, etc. In this study, a total of 65 membrane proteins have been identified. Over eighty percent (80%) of the membrane proteins had unknown functions, many of which may contribute to the antigenic diversity of the virus. Previous studies showed that immunized pigs with the baculovirus expressed hemagglutinin of ASFV were protected against the viral lethal infection ^28^. The results suggested that incorporation of multiple antigens in the vaccine may provide better protection. Therefore, much more efforts are needed to determine the role of membrane proteins in stimulating neutralizing antibodies, and to investigate the neutralization mechanisms and efficiency of the antibodies.

Indels were found to have larger contribution to the genetic diversity of ASFVs than the point mutations. Compared to point mutations, indels could introduce a larger variation to the genome, and cause a more severe damage to the genome structures, which may lead to the death of viruses. Therefore, only few indels were observed in viruses with small genomes, such as influenza viruses and hepatitis B viruses (HBV). However, it was more robust for the indels to occur inside the viruses with large genomes, such as ASFVs and poxviruses ^3,29^, because the viruses with large genomes had lots of repeated elements and duplicated proteins (paralogs). Moreover, indels may provide a more efficient way of survival than the point mutations under the natural selection pressure, since the virus with indels could rapidly change its phenotype ^3,29^, such as antigen, virulence, or ability of replication and transcription. For example, the deletion of some MGF genes in ASFV could reduce the viral replication or virulence, which may help with the viral infection of soft ticks ^3,30^.

Several factors could contribute to the indels and the gene duplications, including replication slippage, retrotransposition, recombination, aneuploidy, polyploidy, etc ^31^. The replication slippage may introduce short indels which were widely observed in ASFV genomes, but it is unlikely to cause large indels. This study has demonstrated that the ectopic homologous recombination ^32^, during which the segments with unequal length were exchanged (Figure 6A), may contribute much to the extensive indels observed in ASFV genomes. As a proof, significant associations were observed between the occurrence of extensive recombination events and the indels. Two factors may facilitate the homologous recombination in ASFVs: firstly, a large amount of clustered repeated elements were observed in ASFV genomes (Figure 5); secondly, all the analyzed ASFVs in this study contained a possible recombinase and DNA topoisomerase, both of which were commonly observed enzymes responsible for homologous recombination. Both of enzymes were very conservative and experienced no recombination, suggesting their important roles in ASFVs. Taken together, the homologous recombination should be the effective strategy of ASFVs to generate the genetic diversity, which further leads to the diverse phenotypes, including antigen, virulence, replication and transcription ability, and the “weapons” of escaping from the host immunity (Figure 6B).

**Figure 6.**
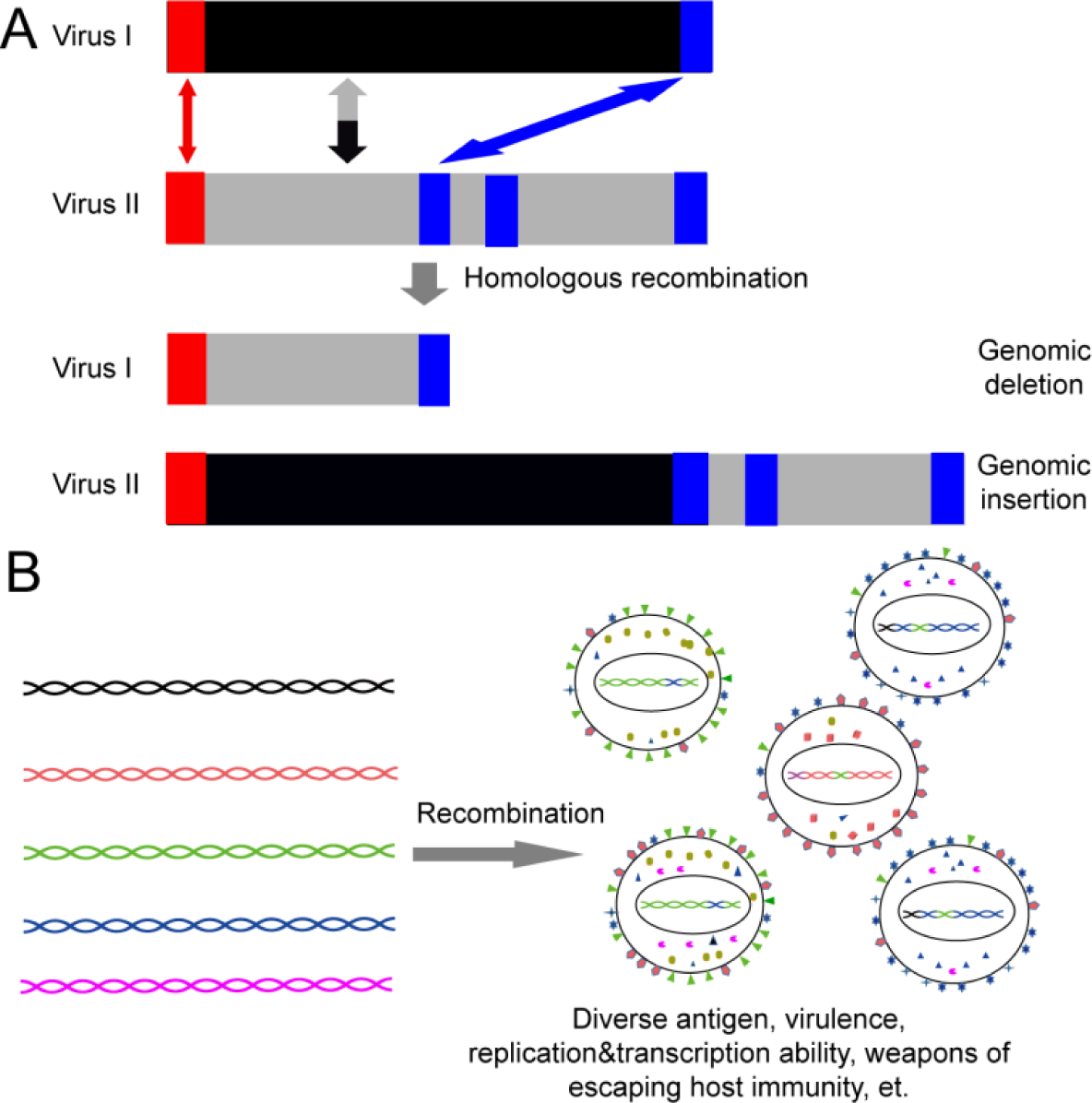
Homologous recombination leads to (A) the indels, and (B) the genetic diversity of ASFVs.

There were some limitations to this study. Firstly, the number of ASFV genomes was limited, which hindered a comprehensive analysis on the evolution of ASFV genomes. Fortunately, the isolates included in this study covered a long time period from 1950 to 2017, and also covered a large area including Africa and Europe, which were the two major areas of the ASFV circulation. Thus the results based on these isolates reflect the genetic diversity of the ASFVs to a large extent. Secondly, the location and size of the indels observed in ASFV genomes may be affected by the sequence alignment algorithm. Two common methods for the alignment of ASFV genomes were used in this study. In both methods, frequent indels were observed, and the indels were demonstrated to be more responsible for the genetic diversity than the point mutations. Thirdly, the proteome of each ASFV was inferred by computational methods. All of the obtained proteins had significant homology to the proteins in the NCBI Protein database, however, most of the proteins in the NCBI Protein database were only predicted without experimental validations. Besides, functions of nearly 70% of ASFV proteins were unknown. Further experimental studies were needed to determine the proteome and the functions in ASFVs ^14,15^. Lastly, the extensively repeated elements in ASFV genomes could facilitate the frequent occurrence of recombination events. However, some of recombination events cannot be detected by the recombination detection method because of the exchange between the genomic segments with small indels. Such kinds of recombination events are difficult to detect. Increasing the sensitivity of the recombination detection method can help detect them, but may also bring false positives. Therefore, the sensitivity and specificity should be balanced in the recombination detection methods.

Overall, this work provided a systematic view of the genetic diversity of ASFVs. Extensive homologous recombination detected in this study may contribute much to the widespread indels observed in ASFV genomes, which further lead to the large genetic diversity of ASFVs. The results on the causes of the diversity of ASFVs would help with the understanding of the evolution of the virus and thus facilitate the prevention and control of ASFVs.

## Materials and Methods

### 1 ASFV genome and alignment

All the ASFV genomic sequences with over 160, 000 bp were obtained from NCBI GenBank database on October 7, 2018 ^33^. After removing the genomic sequences derived from a patent, a total of 36 ASFV genomes were kept in the analysis. The genomic sequences were aligned by MAFFT (version 7.127b) ^34^. To ensure the robustness of the alignment, the traditional tool of CLUSTAL (version 2.1) ^35^ was also used to align these genome sequences.

### 2 ORF prediction

To obtain the proteins encoded by the ASFV genomes, each genome sequence was searched against all the ASFV protein sequences obtained from the NCBI protein database on October 7, 2018, with the help of blastx ^36^. All genomic regions with significant hits (e-value < 0.001) were checked using a Perl script: overlapping regions in the same coding frame were merged to obtain open reading frames (ORFs) as long as possible; regions without start codon or stop codon were extended upstream or downstream to search for the start or stop codon. Then, the genomic regions which had i) significant hit, ii) both sequence identity and query coverage percentage greater than 60%, iii) both start and stop codons, and iv) over 120 bps, were defined as the candidate ORFs. The candidate ORFs were then translated into proteins using a Perl script. The proteins, which were either completely embedded within another protein, or contained less than 40 amino acids due to early termination of translation, were removed.

### 3 Protein grouping

All the inferred proteins of ASFVs were grouped based on sequence homology using OrthoFinder (version 2.2.7) ^37^ with the default parameters. Manual check was conducted to ensure that each protein group contains one type of protein.

### 4 Calculation of the ratio of dN/dS for proteins

The coding sequences of proteins in each protein group were aligned by codon according to the protein sequence alignment using a Perl script. The ratios of dN/dS between pairwise coding sequences were calculated by yn00 in PAML (version 4.1) ^38^. The average of pairwise dN/dS ratios was calculated as the ratio of dN/dS for the protein.

### 5 Alignment of ASFV proteome

An ASFV proteome was defined as all the proteins encoded by the ASFV genome. Because both the plus and minus strands could encode proteins, the proteins in a proteome were separated into plus and minus proteome based on the coding strands. Proteome alignment was conducted separately for the plus and the minus proteomes. Firstly, proteins in each proteome were sorted with the order from the 5’ end to the 3’ end of the genome, based on the coding regions of the proteins. Then, the proteomes were aligned using a dynamic programming algorithm. Manual check was conducted to ensure that there was no mismatch of proteins in the alignment.

### 6 Function inference and classification of ASFV proteins

The name of each protein group was obtained from the names of BLAST best hit of proteins included in the protein group. To infer the function of each protein group, the longest protein sequence in each protein group was selected as the representative of the protein group. InterproScan (version 5) ^39^ was used to infer the function of the representative protein sequence. The TMHMM Server (version 2.0) ^40^ was used to predict whether the representative protein had a trans-membrane helix. Membrane proteins were defined as those who had at least one trans-membrane helixes. The functional classification of the proteins was adapted from Dixon’s ^3^ and Alejo’s ^15^ work.

### 7 Detection of homologous recombination events

RDP (version 4) ^41^ was used to detect the recombination events in the aligned ASFV genomes. Multiple methods in RDP were used. Only the recombination events which were detected by at least two methods were used for further analysis.

### 8 Evolutionary analysis of YqaJ-like viral recombinase domain

All viral protein sequences of the family of Yqaj (YqaJ-like viral recombinase domain, ID: PF09588) were downloaded from the Pfam database ^42^ on November 21, 2018. With the help of blastp, the recombinase domain in ASFV was found to be most similar to that in two giant viruses, Pacmanvirus and Kaumoebavirus, with sequence identities equal to 0.35 and 0.30, respectively.

### 9 Searching for retrotransposon in ASFV genomes

All retrotransposons in the databases of RepBase (Version 23.10) ^43^ and TREP ^44^ were downloaded on November 11, 2018. All ASFV genomes were searched against these retrotransposons using blastn. No hits were obtained under the e-value cutoff of 0.001.

### 10 Phylogenetic tree inference and visualization

Maximum-likelihood phylogenetic trees were inferred using MEGA (version 5.0) ^45^ with the default values of parameters. Bootstrap analysis was conducted with 100 replicates. The phylogenetic tree was visualized using Denscrope (version 2.4) ^46^.

### 11 Statistics analysis

All the statistical analyses were conducted in R (version 3.2.5) ^47^.

## Supporting information

## Acknowledgements

This work was supported by the National Key Plan for Scientific Research and Development of China (2016YFD0500300, 2016YFC1200200 and 2017YFD0500104), the National Natural Science Foundation of China (31500126 and 31671371) and the Chinese Academy of Medical Sciences (2016-I2M-1-005).

