## Supplementary material for "Homologous Recombination as an Evolutionary Force in African Swine Fever Viruses"

**Supplementary Figures**

**Figure S1**. Comparison of genomic differences caused by indels and point mutations. Alignment of ASFV genomes were conducted with CLUSTALW.

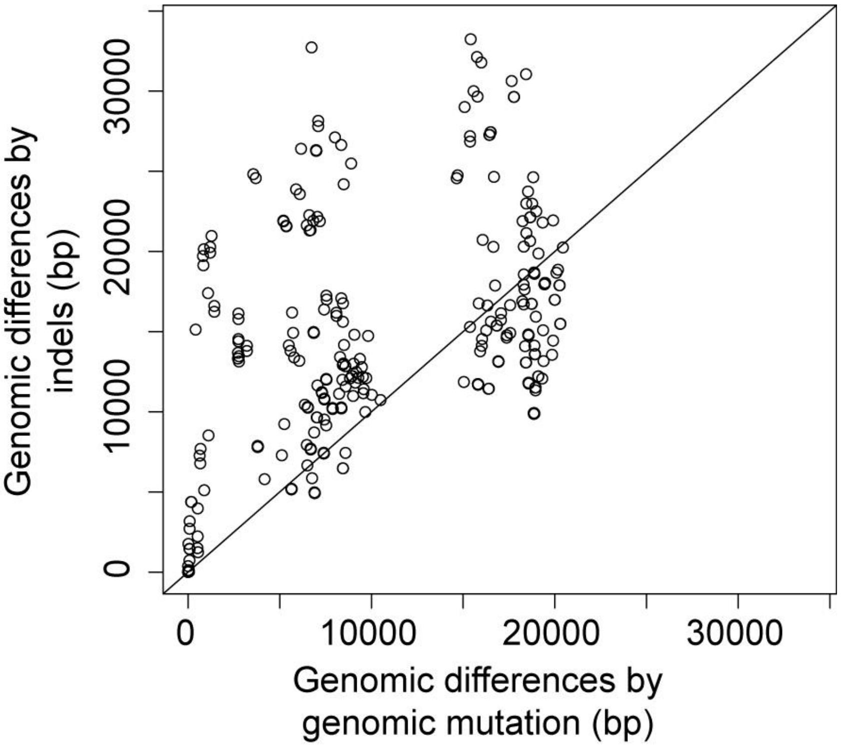


**Figure S2**. Distribution of indel size in ASFV genomes.

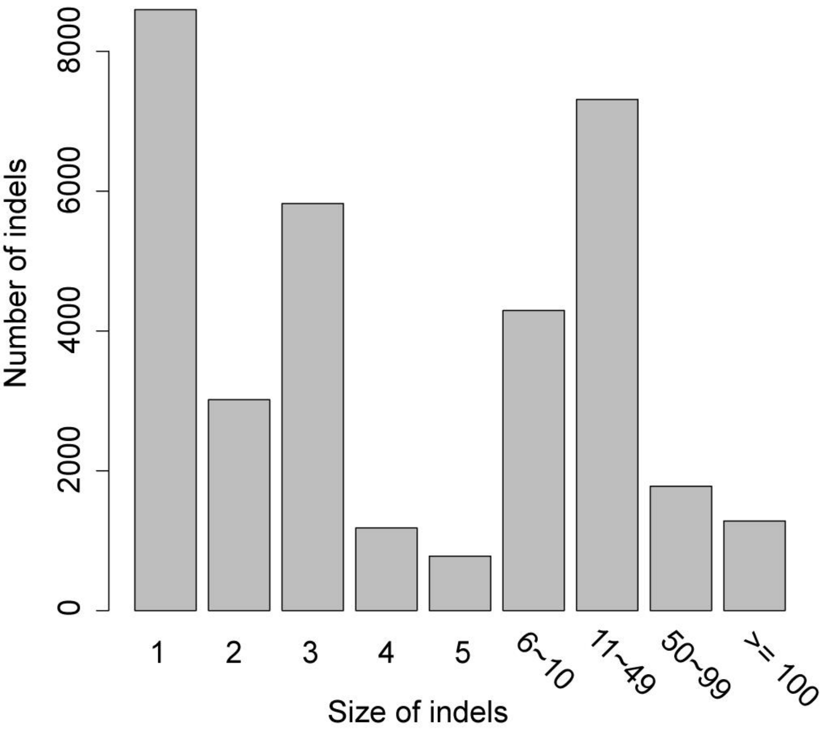

**Figure S3**. The functional composition of protein groups. Numbers in square refer to the number of core protein groups versus that of variable protein groups.

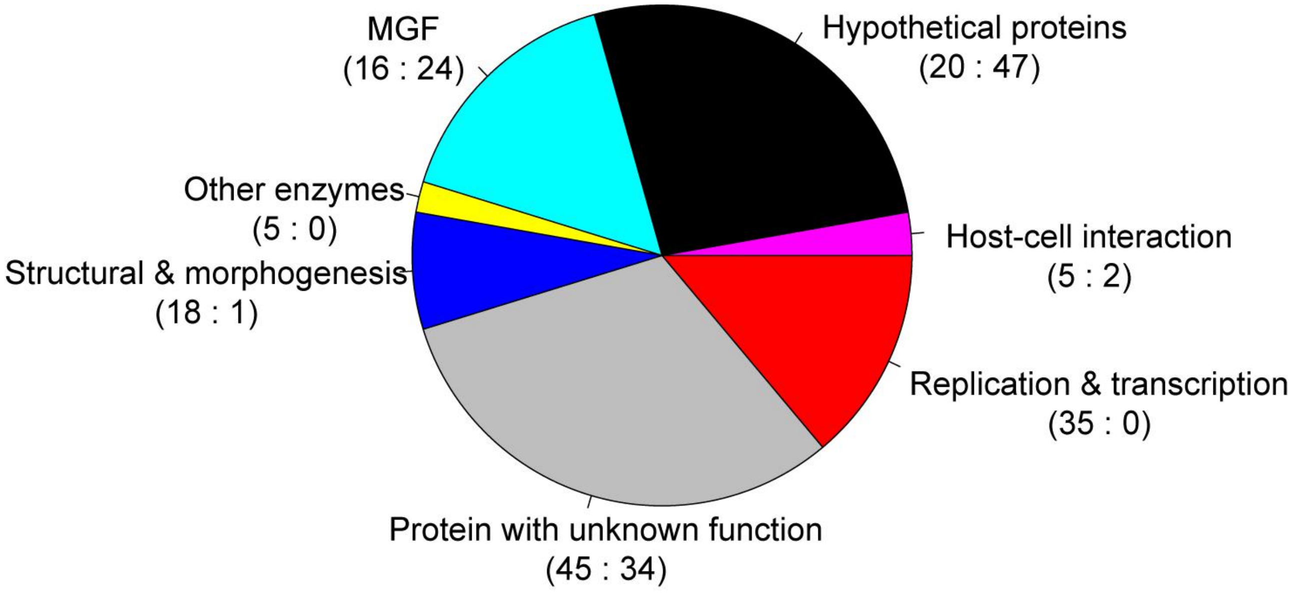


**Figure S4**. The distribution of pairwise protein sequence identities for proteins in each functional class.

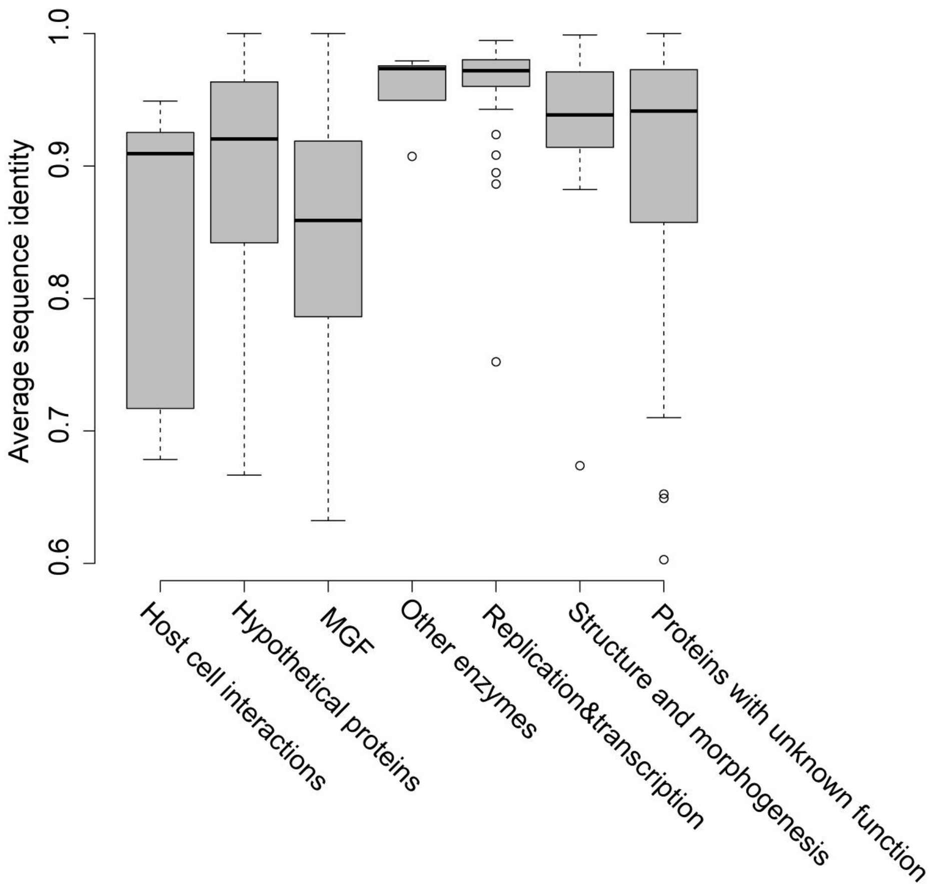

**Figure S5**. Distribution of ratios of dN/dS for proteins in each functional class. For clarity, ratios of dN/dS greater than 3 were truncated to 3.

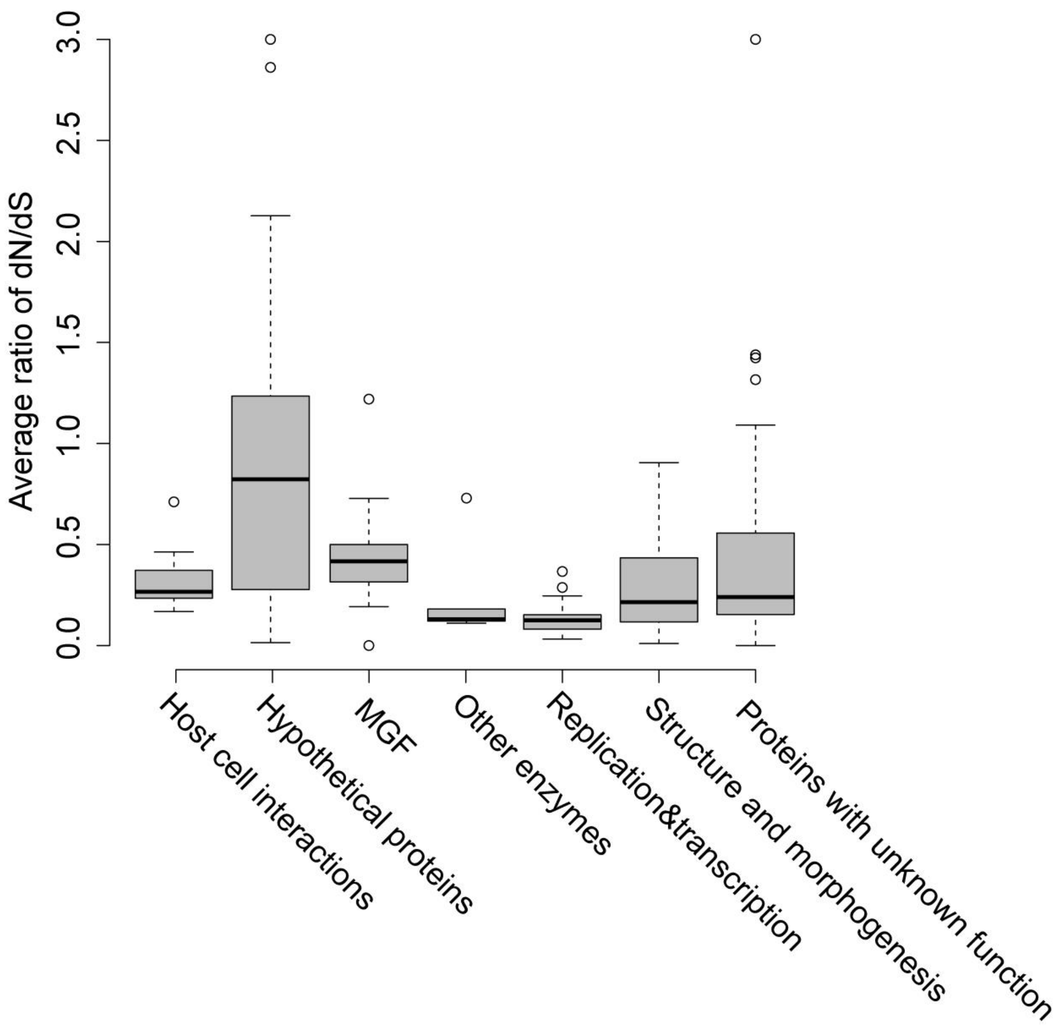

**Figure 6**. The phylogenetic tree of ASFVs and the alignment of their proteomes in plus (left side) and minus strand (right side). Each row refers to the proteome of the virus listed in the phylogenetic tree; each colume refers to one protein group. Protein groups with at least two paralogs in at least one virus were colored, while the others were colored in gray. "White" refers to no protein group in the virus. For clarity, the singletons were ignored in the alignment. The black arrow points to the protein group of p01990-3L.

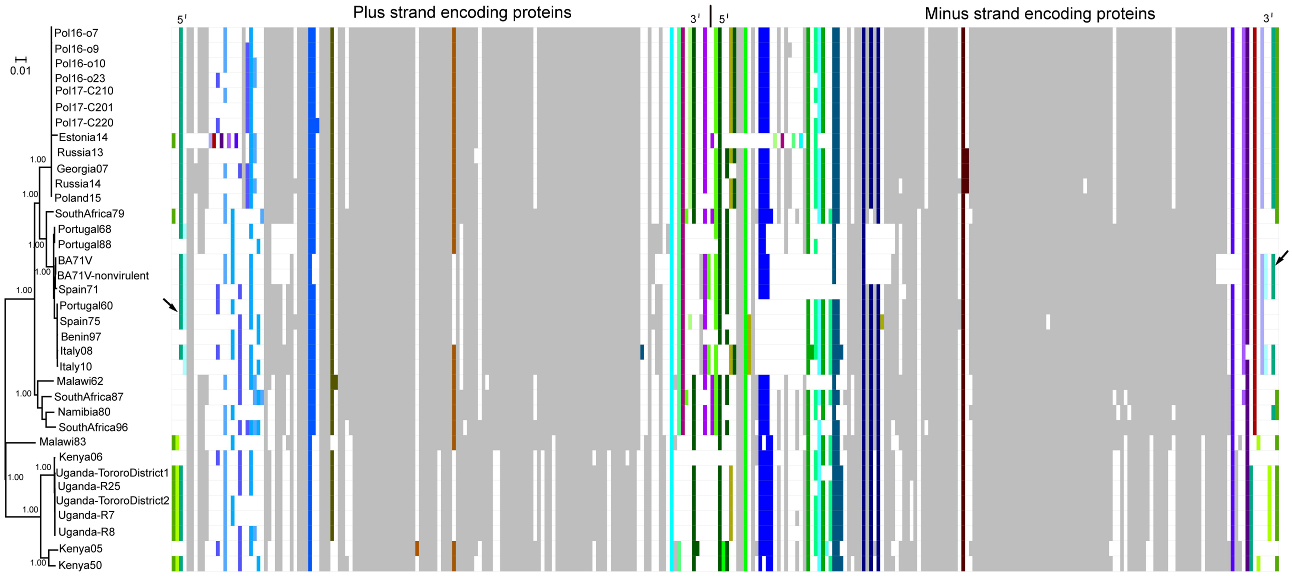

**Figure S7.** Comparison of the ratios of gaps in the recombination regions and other regions. The mean ratio of gap in recombination regions and other regions was 0.23 and 0.03, respectively, while the median ratio of gap in these regions was 0.06, 0, respectively.

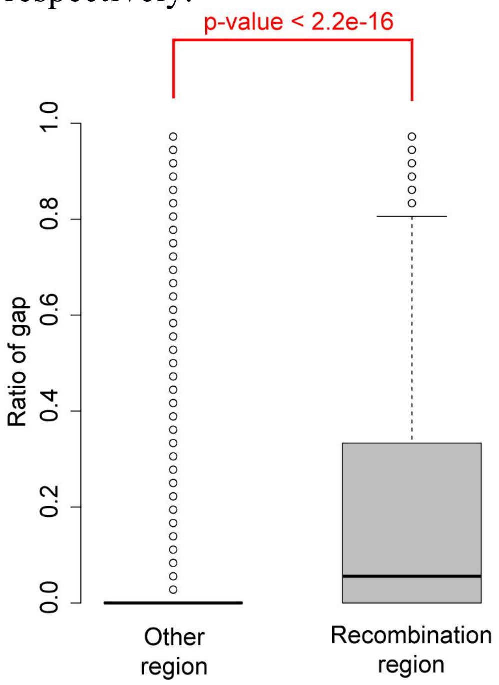


**Figure S8.** Comparison of the number of indels in the recombination region and other regions. The indels greater than 5, 10 and 50 bp were considered here. In all cases, the number of indels in the recombination region is significantly larger than those in other regions. "***", p-value < 0.001.

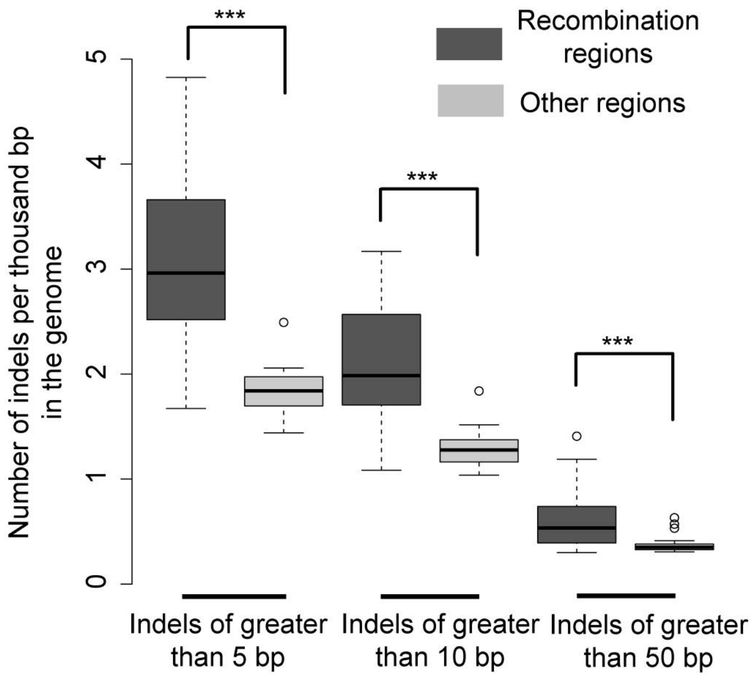

**Figure S9**. The phylogenetic tree of pD345L (A) and P1192R (B).

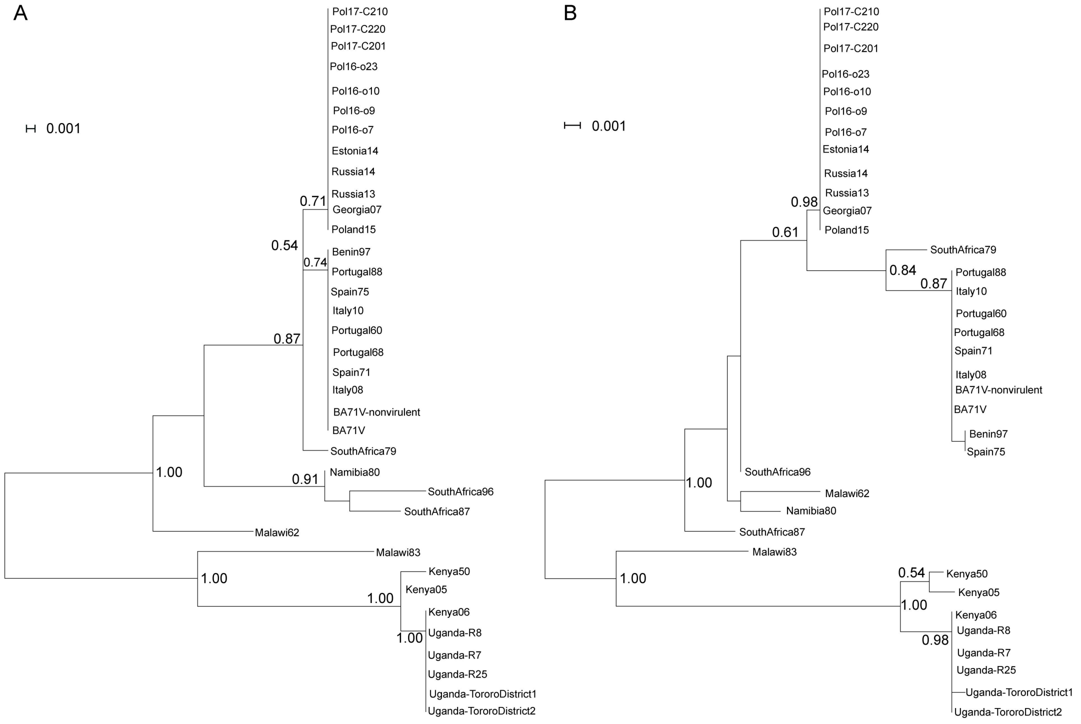


**Figure S10**. The median numbers of repeated elements in ASFV genomes versus the size of repeated elements. For clarity, the natural logarithm of the number of repeated element was used.

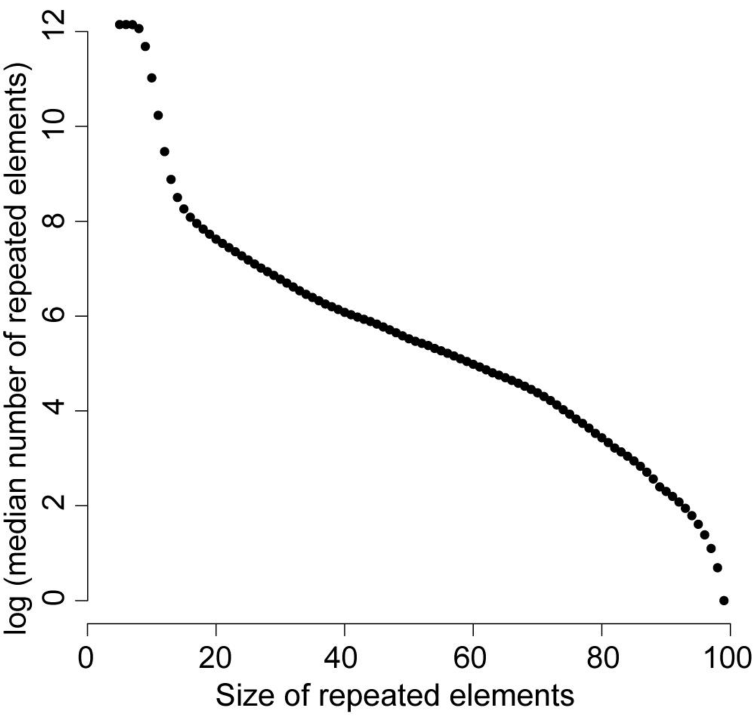
